# *Afrothismia ugandensis* sp. nov. (*Afrothismiaceae*), Critically Endangered and endemic to Budongo Central Forest Reserve, Uganda

**DOI:** 10.1101/2024.04.03.587962

**Authors:** Martin Cheek, Roy E. Gereau, James Kalema

## Abstract

The fully mycotrophic *Afrothismia ugandensis* (Afrothismiaceae), formerly described as *A. winkleri* var. *budongensis* Cowley, is renamed, redescribed and illustrated from the Budongo Forest in Western Uganda. This change in status is supported by eight newly elucidated qualitative morphological diagnostic characters despite the overall similarity with *A. winkleri*, a species restricted to Cameroon and Gabon. *Afrothismia ugandensis* is remarkable in the genus for occurring in semi-deciduous (not evergreen) forest and for having ellipsoid or ovoid (vs. globose) root bulbils. It has only been recorded twice, first in August 1940, and most recently in June 1998, despite targeted searches in recent years. In both 1940 and 1998, only single individuals appear to have been detected. A single site for the species is known with certainty. It is here assessed as Critically Endangered (CR B2ab (iii); D1) using the IUCN 2012 categories and criteria. *Afrothismia ugandensis* is threatened by forest degradation and clearance due to illegal selective small-holder logging for firewood and charcoal, timber and limited agriculture.

## Introduction

Fully mycotrophic heterotrophs, often known as achlorophyllous mycotrophic plants, or saprophytes, are remarkable for lacking all chlorophyll and being completely dependent on fungi for their survival. In continental Africa, individual species or entire genera that are achlorophyllous mycotrophs occur in the families Orchidaceae, Gentianaceae and Burmanniaceae, while all members of Afrothismiaceae, Triuridaceae and Thismiaceae are fully mycotrophic (Cheek & Ndam 1996; Cheek & Williams 1999; Cheek 2006; Cheek et al. 2023a).

Although some earlier authors (Jonker 1938; Maas-van der Kamer 1998) placed *Afrothismia* (Engl.) Schltr. and associated genera as a tribe Thismieae within the family Burmanniaceae sensu lato, molecular phylogenetic data (e.g. Merckx et al. 2006) strongly indicate that Thismiaceae are best placed as a separate family (Cheek et al. 2018a). Further, subsequent relatively well-sampled analyses place Afrothismiaceae as sister to Taccaceae +Thismiaceae, in a different subclade of Dioscoreales from Burmanniaceae sensu stricto (Merckx et al. 2009; Lin et al. 2022). The families are separated from each other by numerous morphological characters and Afrothismiaceae was formally recognised recently (Cheek et al. 2023a).

Afrothismiaceae, with a single genus, *Afrothismia*, are confined to tropical continental Africa (not a single species is known from Madagascar, the Comores nor the Mascarenes). *Afrothismia* differs from the genera of Thismiaceae by the annulus inserted deep inside the perianth tube; stamens inserted below the annulus; anthers usually adnate to stigma; rhizomes with clusters of usually spherical tubers and being confined to tropical equatorial central and eastern Africa (Cheek et al. 2018a). Sixteen species have been formally described (now 17, with this paper), but there appear to be seven additional undescribed species (Cheek et al. 2023a).

The genus *Afrothismia* was erected by Schlechter (1906), based on *Thismia* sect. *Afrothismia* Engl (1905). He transferred to it *Thismia winkleri* Engl. (Engler 1905) and described a second species, *A. pachyantha* Schltr., that he had collected in the then German colony Kamerun, now Cameroon. The range of the genus was formally extended to E. Africa by Cowley (1988), with *A. insignis* Cowley from Tanzania and *Afrothismia winkleri* var. *budongensis* Cowley from Uganda. By 2003, the original two species known for the genus had increased in number to four, including the Cameroonian species, *A. gesnerioides* H.Maas (Maas-van der Kamer 2003).

In the ensuing 15 years, the number of formally described new species quadrupled to 16, with discoveries from Gabon (Dauby et al. 2008), Kenya (Cheek 2004), Malawi (Cheek 2009) and Tanzania (Cheek & Jannerup 2006); but the largest number of discoveries by far has been made in Cameroon. Here, eight new species of *Afrothismia* have been published, one extending to Nigeria (Franke 2004; Franke et al. 2004; Sainge & Franke 2005; Sainge et al. 2005; Cheek 2007; Sainge et al. 2013; Cheek et al. 2019). Most of the Cameroonian species fall within the Cross-Sanaga interval (Cheek *et al*. 2001), which holds the highest flowering plant species (Barthlott et al. 1996) and generic (Dagallier et al. 2020) diversity per degree square in Tropical Africa, and which is the site of many new discoveries, including genera new to science (Litt & Cheek 2002; Cheek et al. 2003; Cheek et al. 2018b). Many of the Cameroonian *Afrothismia* species feature in the Red Data Book of Cameroon Plants (Onana & Cheek 2011) and most species of the genus are Critically Endangered (Cheek et al. 2023a) including the type species, *A. winkleri*, until now considered to include the Ugandan taxon (IUCN SSC East African Plants Red List Authority 2013). One Cameroonian species is considered extinct after targeted searches (Cheek et al. 2019), while several others have not been recorded alive in several decades and so may also be extinct, e.g. *A. zambesiaca* Cheek (Cheek 2009).

Although species niche modelling has indicated that *Afrothismia* might occur in West Africa (Sainge et al. 2017), recent targeted expert surveys there have failed to find any species of the genus, although they have produced other new species of fully mycotrophic Dioscoreales (Cheek & van der Burgt 2010; Cheek et al. 2023b).

Lacking any green tissue, *Afrothismia* spp. depend on vascular arbuscular mycorrhizal fungi for sustenance. The genus *Rhizophagus* P.A. Dang (Glomerales, Glomeromycota) has been implicated as the fungal partner of the genus (Franke et al. 2006). Delayed co-speciation between *Afrothismia* and the fungal partner has been demonstrated (Merckx 2008; Merckx & Bidartondo 2008). However, the autotrophic partners of these fungi remain unknown.

The new species of *Afrothismia* reported in this paper is a result of the Uganda Tropical Important Plant Areas programme (TIPAs, see Darbyshire et al. 2017; Richards et al. 2024).

Cowley (1988) recognised the Ugandan taxon as different at varietal rank from *Afrothismia winkleri*. This was based on quantitative characters and a difference in ovary colour (Cowley 1988). In this paper we show that there are eight additional, qualitative diagnostic characters supporting separation of the Ugandan material from *Afrothismia winkleri*. These are more than adequate for full species recognition, and we here formally describe the available material as *Afrothismia ugandensis*.

## Materials & methods

Nomenclatural changes were made according to the *Code* (Turland et al. 2018). Names of species and authors follow IPNI (continuously updated). Herbarium material was examined with a Leica Wild M8 dissecting binocular microscope fitted with an eyepiece graticule measuring in units of 0.025 mm at maximum magnification. The drawing was made with the same equipment with a Leica 308700 camera lucida attachment. The following herbaria were inspected for specimens: B, BM, EA, K, MHU, SRGH, YA, WAG. Herbarium codes follow Index Herbariorum (Thiers, continuously updated).

The use of technical terms follows Beentje & Cheek (2003) and the format of the description follows those in other papers describing new species in *Afrothismia*, e.g. Cheek et al. (2004, Cheek & Jannerup 2006; Cheek et al. 2019). The specimens cited have all been seen by one or other of the co-authors. The conservation assessment follows the IUCN (2012) categories and criteria.

## Results

The species described in this paper as *Afrothismia ugandensis* Cheek was first encountered, collected and preserved in spirit in the Budongo Forest of Uganda in August 1940 by Eggeling (*Eggeling* 4041, K) who made a watercolour sketch of it (Fig. 1). The specimen was initially determined at Kew as *A. winkleri*, but included in the Flora of Tropical East Africa Burmanniaceae account as *A. winkleri* var. *budongensis* Cowley (Cowley 1988). The taxon was differentiated from *A. winkleri* based on quantitative characters and a difference in colour of the ovary (Cowley 1988).

**Fig. 1.**
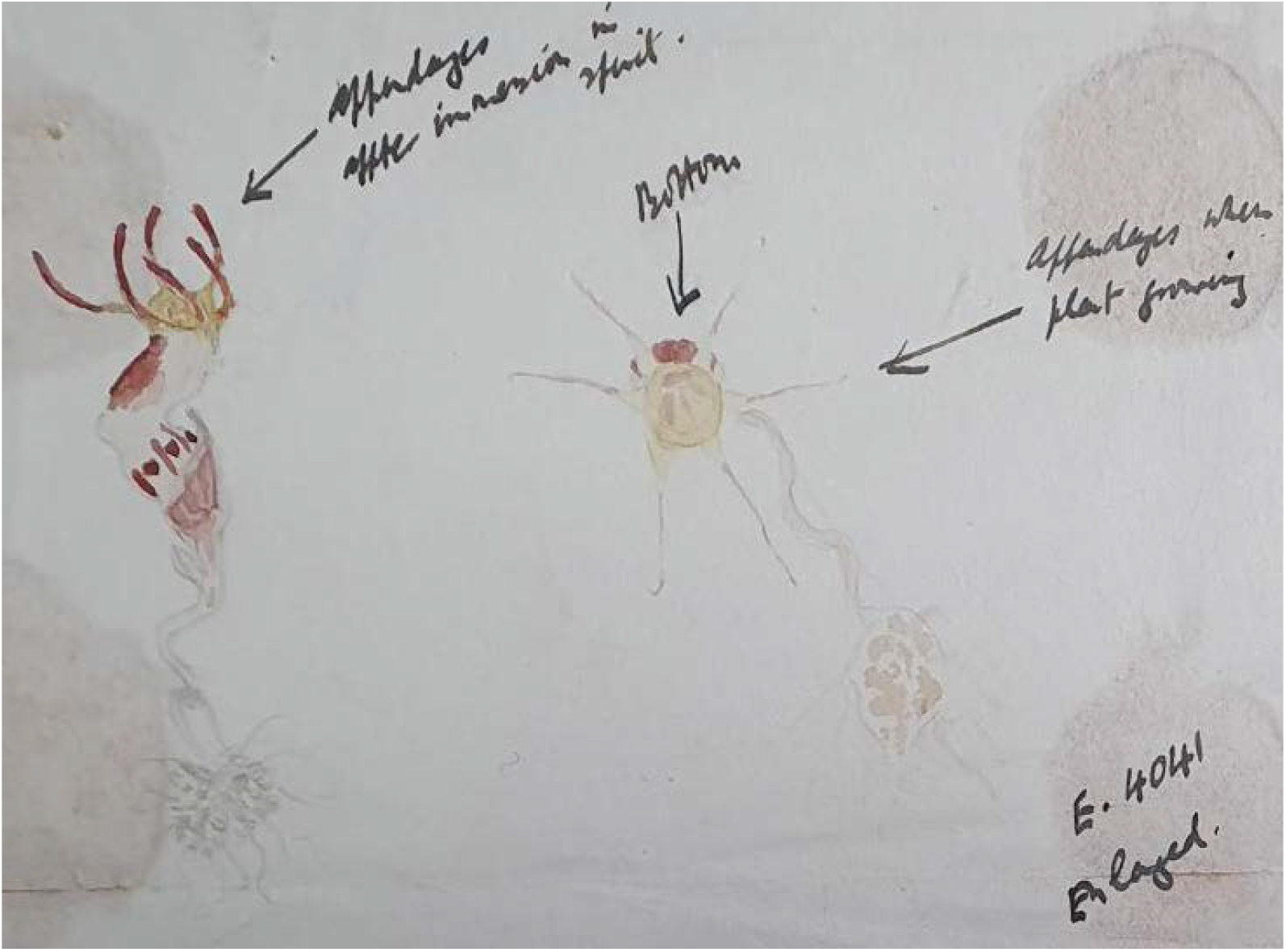
*Afrothismia ugandensis* Cheek. Water coloured field sketch of *Eggeling* 4041 (K spirit collection number 3669): left, flowering plant, showing flower in side view annotated presumably by Eggeling “appendages after immersion in spirit” and right, plan view “appendages when plant growing”. Mounted on the herbarium sheet cross-referenced to the spirit (and only) surviving specimen. Drawn by W. J. EGGELING.

However, the two taxa are separated geographically by c. 2300 km and morphologically using the eight additional qualitative characters indicated in Table 1 and in the diagnosis, more than adequate for species level rather than varietal status. *Afrothismia ugandensis* is remarkable for being the only published species of the genus from Uganda and the westernmost member of the family in East Africa (Cheek et al. 2023a).

**Table 1.** The characters separating *Afrothismia ugandensis* from *Afrothismia winkleri*. Data for the second species from Engler (1905).

|  | *Afrothismia ugandensis* | *Afrothismia winkleri* |
| --- | --- | --- |
| Tepal lobes | Base hastate, with flanking pair of subsidiary triangular lobes | Base entire, lacking any lobes |
| Corona surface | Densely papillate | Smooth |
| Annulus surface | Densely puberulent | Glabrous |
| Annulus shape | Entire, constant in width, continuous | Divided into 6 broadly triangular lobes |
| Anther connective distal appendage attached to stigma? | Absent, stamen not attached to stigma | Present, ovate, attached to stigma |
| Style surface | Glabrous | Densely puberulent |
| Outer (abaxial) stigma surface | Glabrous | Sparsely puberulent |
| Stigma shape and lobing | Divided nearly to the base into three triangular/oblong lobes, each lobe itself bifid | Shallow bowl minutely 6-lobed at margin |
| Geographic range | Uganda | Cameroon and Gabon |

### Afrothismia ugandensis Cheek sp. nov. — Fig. 1, 2

Type: Uganda, Budongo Forest, “Tiny saprophyte 1½ to 2 inches high, growing among dead leaves in dense shade, on the floor of the Budongo Forest, Uganda. Stem and scale leaves pale brown. Perianth with 6 long filiform appendages (straight in life). Perianth partly yellow, partly claret, partly colourless and more or less transparent, as shown in the attached sketch.” fl. Aug 1940, *Eggeling* 4041 (holo. K spirit collection number 3669, barcode K000593368).

**Fig. 2.**
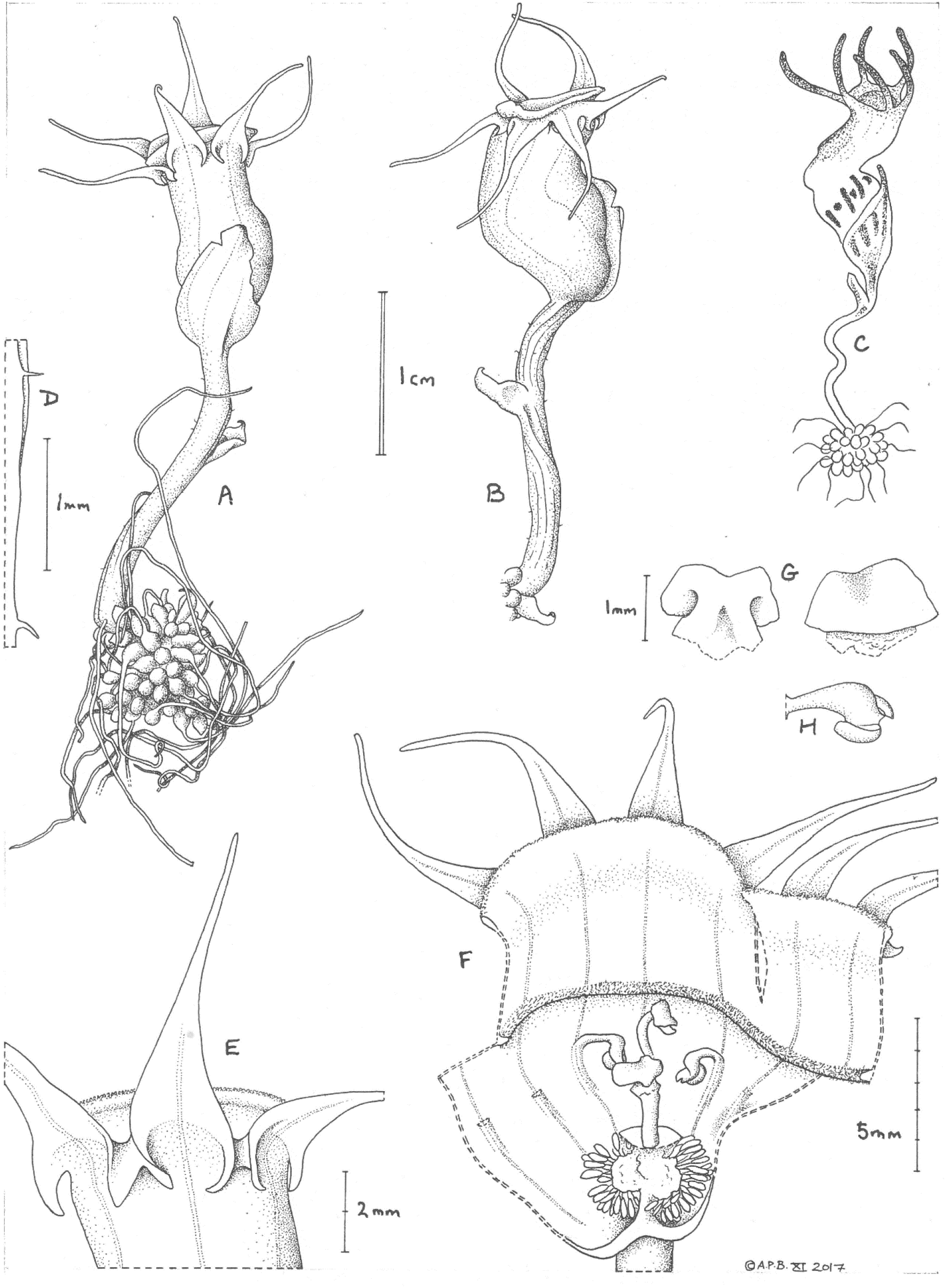
*Afrothismia ugandensis* Cheek. **A**. Habit, whole plant, showing flower in dorsal view; **B**. as A, but showing flower in side view; **C** copy of habit painting by Eggeling (no scale bar was given) presumably of the plant in A & B; **D** detail of stem showing hairs; **E** upper perianth tube showing connation of the tepal lobe bases; **F** dissected damaged flower (only 3 of 6 stamens remain intact); **G** stigma lobe, lower surface (left) and upper (right); **H** stamen viewed in situ. **A-H** from *Eggeling* 4041 (K, spirit collection number 3669). Drawn by ANDREW BROWN.

*Afrothismia winkleri* (Engler) Schltr. var *budongensis* Cowley (Cowley 1988: 7) *Achlorophyllous mycotrophic herb* with only the flower emerging above the leaf-litter. *Bulbil cluster* completely covering the short, naked (scale leaves absent) rhizome, c. 9 – 10 × 8 mm, bulbils ellipsoid or ovoid, each c. 1 – 2.5 × 1 mm, with an apical rootlet up to 3.5 cm long, 0.2 mm diam. (Fig. 2A). *Stem* (peduncle), succulent, concealed in substrate, vertical, unbranched, terete, with 5 – 6 longitudinal ridges, c. 20 – 30 mm long, 2 mm diam., hairs very sparse, simple, patent, 0.2 mm long; scale-leaf (bract) single, inserted approx. midlength, ovate-triangular, c. 0.4 × 0.2 mm, axillary bud globose, internodes c.10 mm long. *Inflorescence* 1-flowered; flower-subtending bract sheathing the base of the flower, translucent when live (field painting), ovate, 7 – 8 × c. 5.5 mm, apex acuminate (Fig. 2C). Flowers slightly asymmetric, c. 1.5 cm long and 2 cm diam. including the perianth lobes (preserved material). *Perianth tube* directed vertically, L–shaped, the axis of the lower perianth tube angled at c. 30 – 40 degrees away from the vertical, the upper tube angled from the lower tube at 50 – 60 degrees in the opposite direction (towards the vertical), tube c. 8 mm long, 5 mm wide at midpoint (at junction of lower and upper tubes). Lower (proximal) tube funnel-shaped, c. 5 mm wide at junction with ovary, widening to c. 6.5 mm wide before constricting to c. 5 mm wide at junction with upper tube, outer surface translucent or white with 6 longitudinal purple lines, each separated by a purple spot on the lower tube and ovary. Upper tube cylindrical, c. 4 × 5 mm, translucent or white with a purple patch in the distal dorsal half, glabrous and lacking projections. Inner surface of perianth tube with an internal flange (annulus) at junction of lower and upper tube, above the insertion of the staminal filaments, flange continuous, unlobed, projecting into the tube c. 0.3 mm, 0.3 mm thick at the base, densely hairy, hairs patent, simple, 0.1 mm long; inner perianth surface otherwise glabrous and lacking ornamentation. Corona yellow, slightly funnel-shaped, projecting 1 – 1.25 mm above the insertion of the perianth lobes, the mouth orbicular in plan view, c. 7 mm diam., the rim thickened, c. 0.25 mm diam., minutely papillose (Fig. 2F). *Tepals* six, monomorphic (equal), patent in life (field painting), (± forward-directed in spirit preserved material), purple, narrowly triangular, flattened, 8 – 9 × c. 2 mm wide at base, tapering gradually to the minutely rounded apex; basal lobes narrowly triangular, c. 1.5 × 0.3 – 0.5 mm, reflexed, curved under the main lobe and connate with the basal lobes of adjoining tepals (Fig. 2E), unornamented, glabrous. *Stamens* six, staminal filaments joined to perianth tube for c. 2.5 mm, appearing as purple lines on exterior tube surface, distal part free, terete 1.8 – 2 mm long, c. 0.25 mm diam., arching inward 180 degrees towards the stigma, distal part swollen abaxially, convex, c. 0.5 mm wide, confluent with the flat expanded connective, together shaped like a duck’s head (Fig. 2F); anthers ellipsoid, c. 0.8 × 0.6 mm, the 2 thecae c. 0.1 – 0.2 mm wide, separated by the connective (Fig. 2H); connective c. 0.5 × 0.5 mm, not extended as a distal appendage, nor adhering to the stigma surface (Fig. 2F). *Ovary* cup shaped, c. 3 mm long, c. 5 mm wide at junction with perianth; placentation axile (placenta attached at base and apex), the ovules inserted on a globose placenta held on a stalk c. 1 mm long (Fig. 2F); ovules ellipsoid c. 1.25 mm long, funicle c. 1 mm long. Style cylindrical, c. 1.5 mm long, 0.5 mm diam. glabrous and unornamented; stigma 3-lobed nearly to the base, lobes flat, patent, smooth, glabrous, oblong triangular in outline, c. 1 × 2 mm, each bilobed, the subsidiary lobes lateral, reflexed (Fig. 2G). *Fruit and seed* not seen.

### RECOGNITION

*Afrothismia ugandensis* Cheek differs from *Afrothismia winkleri* (Engler) Schltr. in the tepal lobes appearing hastate with a pair of subsidiary triangular lobes (vs. no lobes), the corona rim densely papillate (vs. glabrous), the annulus entire, continuous, unlobed, densely puberulent (vs. lobed, glabrous), the style and abaxial stigma glabrous (vs. puberulent), the stigmas deeply 3-lobed, the lobes bifid (vs. stigma forming a shallow bowl with 6 equal, minute lobes)

### DISTRIBUTION & ECOLOGY

semideciduous submontane forest in Budongo Central Forest Reserve; 1040 – 1100 m above sea-level. No additional associated achlorophyllous species are recorded as co-occurring. Associated taxa (collected with the paratype under ATBP numbers 635 to 655) *Caloncoba crepiniana* (De Wild. & T. Durand) Gilg (Achariaceae), *Acalypha neptunica* Müll. Arg. var. *neptunica* (Euphorbiaceae), *Ritchiea albersii* Gilg (Capparaceae), *Strombosia* Blume (Olacaceae), *Elatostema monticola* Hook. f. (Urticaceae), *Peperomia fernandopoiana* C. DC. (Piperaceae), *Brachystephanus africanus* S. Moore and *Brillantaisia nitens* Lindau (Acanthacaeae), *Celtis gomphophylla* Baker (Cannabaceae), *Geophila uniflora* Hiern and *Hymenocoleus hirsutus* (Benth.) Robbr. (Rubiaceae), *Microlepia speluncae* (L.) T. Moore (Dennstaedtiaceae), *Cyclosorus interruptus* (Willd.) H. Itô (Thelypteridaceae), *Campylospermum densiflorum* (De Wild. & T. Durand) Farron (Ochnaceae), *Afromorus mesozygia* (Stapf) E.M. Gardner (Moraceae), *Trichilia rubescens* Oliv. (Meliaceae), *Pellaea dura* (Willd.) Hook. (Pteridaceae), *Heterotis rotundifolia* (Sm.) Jacq.-Fél. (Melastomataceae), *Grewia flavescens* Juss. (Grewiaceae/Malvaceae Grewoideae).

### ADDITIONAL MATERIAL

Uganda (U2); Masindi District; Budongo Forest Reserve, Nyakafunjo Nature Reserve, unlogged forest, 01°42’40“N, 031°31’32“E, 1040 m elev., fl. 24 June 1998, ATBP (African Tropical Biodiversity Programme) 653. Collector S.M. Maishanu (Tropicos specimen ID 1282593 specimen missing, see notes below). “Budongo Forest Reserve, Nyakafunjo Nature Reserve, unlogged forest. Rare saprophyte with a deep floral tube. Leaflets with root swelling. Tube mouth bright yellow with purple patch on upper part of floral tube. Base of floral tube marked with purple streaks.”

This gathering was made by S.M. Maishanu (Saidu Muhammed Maishanu, most recently at the Sokoto Energy Research Centre, Usmanu Danfodiyo University, Sokoto, Nigeria) as part of a training exercise run by the ATBP (https://tropical-biology.org/conservation-projects/) in Budongo. The second author, RG, was with Maishanu at the time and assisted writing the field notes, and RG also searched at that time without success for a second individual of the *Afrothismia*, walking 200 m in each direction searching in the dense herbaceous vegetation along the forest trail. The find was not the result of a targeted search, in fact there was a delay before the specimen was identified.

The specimen was identified by RG in 2000 from memory and label notes. Like all unicate specimens from the ATBP collections, lhe single specimen was left at MHU. It has not been relocated. There is no doubt about the identification since the description of the flower is a perfect match with that of the type specimen (MC pers. obs.).

Some effort has since been made to find more material but with no success yet. During the recent fieldwork expedition to Budongo Central Forest Reserve, the Makerere TIPAs team consisting of four botanists led by JK was joined by two experienced field station parabotanists. These conducted surveys of the forest in mid-October 2023, which was a wet part of the year (ideal for finding *Afrothismiai* in flower), and searched for IPA trigger species, including *Afrothismia*. The survey covered the area surrounding the Field Station within the Nyakafunjo (N3) compartment where the 1998 collection of *Afrothismia ugandensis* was made. This is the forest tract considered by the managers to be the most intact (undisturbed), the Strict Nature Reserve (SNR). The forest canopy was indeed thick and closed, while the forest floor was generally open, with sparse ground vegetation cover. The survey was extended to cover areas beyond the SNR up to c. 3.5 km from the station, and lasted four days, but did not yield any sign of *Afrothismia*. It is to be hoped that the specimen will be re-found one day, with more targeted surveys particularly timing the June-August window.

### CONSERVATION STATUS

Only two collections, each of a single individual, made nearly 60 years apart (Aug. 1940 and June 1998), are known of *Afrothismia ugandensis*, both from the Budongo forest. The earliest collection has no indication of where within the forest it was collected, while the second has geographical coordinates and detailed notes. Using the precautionary principle, therefore, it cannot be ruled out that the species is only known at the single, precisely recorded site. Many other species of the genus are known from a single collection at single sites within larger forest areas (see introduction references). The area of occupancy of *Afrothismia ugandensis* is estimated as 4 km^2^ using the IUCN required 2 km × 2 km cell size and extent of occurrence as slightly larger as required by IUCN (2012) and IUCN Standards and Petitions Committee (2024).

This site is probably one of the most sampled sites for plants in Budongo since it is only 2.45 km from the field station and along a trail. Although the plants are small, and the flower held at ground level, they are not completely inconspicuous due to the yellow and purple flower colour. Studies of *Afrothismia* species in the field in Cameroon (Cheek 2006; Cheek et al. 2004) have found that flowering can be continuous over several weeks or even months at sites but that species can be absent from apparently suitable habitat despite targeted searches (e.g. Cheek et al. 2000, 2010, 2011; Harvey et al. 2004, 2010). The few collections (and only two individuals recorded) at Budongo suggests that the species is genuinely highly range restricted and infrequent. The sole recorded site is close to the southern boundary of the forest where there appear to be fronts of forest clearance approaching about 800 m to the SW and 1.45 km to the SE (Google Earth Pro imagery dated 25 June 2017, Maxar Technologies). Although the forest in Nyakafunjo Nature Reserve was said to be unlogged, not all species recorded at the site are characteristic of deep, closed forest: e.g., *Grewia platyclada, Afromorus mesozygia, Celtis gomphophylla, Ritchiea albersii* (RG pers. obs.).

Budongo Forest and many of its species are currently under immense threat from many sources: population pressure from immigration due to wars and civil unrest in neighbouring countries means cropland is scarce and the forest edges are encroached with burning and clearing; illegal pitsawing of the trees for fuelwood, charcoal and timber is well-organised; agribusiness employs outreach farmers who extend cultivation beyond forest boundaries (Budongo Field Station, continuously updated). Budongo forest is faced with the threat of illegal tree felling, qualifying *Afrothismia ugandensis* to be from a single location whose habitat quality is declining. With only two individuals known (well below the threshold of 50), the species also satisfies the requirements for the Critically Endangered category under Criterion D. The conservation status of *Afrothismia ugandensis* is therefore assessed provisionally here as Critically Endangered (CR B2ab(iii); D1) using the IUCN 2012 categories and criteria.

### ETYMOLOGY

Named for Uganda to which it is both unique (endemic) and the only known species of the genus in the country.

### NOTES

*Afrothismia ugandensis* is remarkable in the genus for occurring in semi-deciduous (vs. evergreen) forest and for having ellipsoid or ovoid (vs. globose) root bulbils. The non-attachment of the stamens to the stigma is also unique in the genus, but this observation comes with the caveat that it is based on material damaged by a previous dissection (e.g. 3 of the 6 stamens are missing) and needs to be confirmed if and when new material becomes available to be sure that it is not an artefact of damage.

That *Afrothismia ugandensis* has been considered a variety of *A. winkleri* is unsurprising given their overall similarities and they may well share a recent common ancestor. The Budongo forest is considered the easternmost part of the Congolian phytogeographic domain (White 1983). Until recent geological times, the forest habitat of *Afrothismia* extended continuously across the Congo basin to the Cameroon home of *A. winkleri*, but this forest was only intermittently present in much of the Pleistocene, being replaced with grasslands in colder and drier periods. It is conceivable that the common ancestor of the two taxa once extended across the basin in wetter periods, but that separation in dry periods resulted in the two species diverging. *Afrothismia ugandensis* has little similarity with the Kenyan *A. baerae* Cheek where the lower and upper perianth tube are share the same vertical axis, and the corona mainly occludes the mouth. The Ugandan species shares more similarities with the Tanzanian Eastern Arc species *A. mhoroana* Cheek and *A. insignis* but lacks the ± globose lower tube axis and verrucate perianth of those species.

The collector and illustrator of the type and only specimen seen for this paper, Joe (William Julius) Eggeling (1909 to 1994) joined the British Colonial Forestry Service in 1931 and was first posted to Uganda where his inventory and management plan of the Budongo Forest in Bunyoro was considered a masterpiece in tropical forestry. Thirty-one plant names on IPNI (continuously updated) are named in his honour including the genus *Eggelingia* Summerh. (Orchidaceae). These are mainly Ugandan and Tanzanian species that he brought to light from among the c. 3800 herbarium specimens he collected and deposited at BM and K. He became Chief Conservator of Ugandan forests in 1945 and in his retirement in his native Scotland led many conservation initiatives that led to the creation of areas protected for biodiversity conservation (Dalyell 1994; Wikipedia continuously updated).

Budongo Central Forest Reserve covers an area of 822 km^2^. “It qualifies as an Important Plant Area under criterion A. Sub-criterion A(i) is triggered by the presence of one Critically Endangered, three Endangered and 16 Vulnerable species, while A(iii) is triggered by the presence of highly restricted endemics, *Afrothismia winkleri* var. *budongensis* (DD) and *Uvariopsis* sp. nov. 1 ‘Uganda’” (IPA data sheet, Richards & Darbyshire 2024). Further detailed and uptodate botanical data characterising the Budongo Central Forest Reserve is given in this reference.

Many important aspects of the biology of this fascinating genus remain imperfectly or completely unknown, such as pollination biology (only observed in one species), microsporogenesis, cytology, and seed dispersal. Budongo Forest offers the possibility of filling in these gaps in knowledge for the genus in general, as well as for *Afrothismia ugandensis* in particular.

## Discussion

The case of *Afrothismia ugandensis*, formerly included in the apparently greatly disjunct Cameroon/Gabon species *A. winkleri* (albeit recognised as varietally different), suggests that it might repay effort to reexamine other such cases of apparently far disjunct species with this range. A recent re-assessment of another such disjunct species (western Uganda and Cameroon) species, *Keetia purseglovei* Bridson (Rubiaceae), showed that the Cameroon material was a separate species from the Ugandan (Cheek & Onana 2024). A further example of an apparently widespread and Cameroon/Ugandan disjunct taxon, *Dovyalis spinosissima* Gilg (Salicaceae) also led to the recognition of distinct species in both countries after further taxonomic study (Cheek & Ngolan 2007). This taxonomic work matters because range restricted and nationally endemic species can otherwise be overlooked and not properly prioritised for conservation, increasing their risk of global extinction. Until species are documented, described and known to science, it is difficult to assess them for their IUCN Red List conservation status, and therefore the possibility of conserving them is reduced (Cheek *et al*. 2020). An additional example of a narrowly endemic threatened species in Uganda is *Encephalartos whitelockii* P.J. Hurter (Zamiaceae), which is also Critically Endangered (Kalema 2010). It is endemic to Mpanga Gorge, where at least one other potentially threatened species is in the course of elucidation (e.g. Mucaleque et al. 2024).

Perhaps the highest priority for such range restricted threatened species is to conserve them in their natural habitats (e.g., by including them in Important Plant Areas, Darbyshire *et al*. 2017) and to develop species conservation action plans to improve the likelihood of their survival (e.g. Couch *et al*. 2022). Both Budongo and Mpanga Gorge are already evidenced as IPAs. Conservation *in situ* is crucial for fully mycotrophic species such as *Afrothismia ugandensis* since *ex situ* conservation is currently impossible when the autotrophic partner(s) of the relationship is unknown. Global extinction is a high possibility with highly range restricted, infrequent, single site endemic species such as *Afrothismia ugandensis*. Effort is needed if it is not to follow other African fully mycotrophic species that are already well documented as extinct (Onana & Cheek 2011; Cheek & Onana 2011; Cheek et al. 2018a, 2019)

## Acknowledgements

We thank Melissa Bavington for her care of the spirit collection at K, including the specimen on which this paper is based.

We thank two anonymous reviewers for constructive comments on an earlier version of this manuscript.

The authors declare they have no conflict of interest.

